# Sleep and emotional memory: translating neural response at encoding to memory accuracy in men and women with and without PTSD

**DOI:** 10.64898/2026.03.27.714805

**Authors:** Anthony C. Santistevan, Nikhilesh Natraj, Leslie M. Yack, Kim L. Felmingham, Steven H. Woodward, Daniel H. Mathalon, Thomas C. Neylan, Anne Richards

## Abstract

**Background:** Growing evidence suggests that sleep plays an important role in PTSD outcomes, potentially due to its influence on emotional memory consolidation, though these mechanisms remain unknown. This study sought to test the hypotheses that sleep neurophysiology, PTSD status, and sex moderates the degree to which the late positive potential (LPP) mediates memory accuracy for affective visual stimuli.

**Methods:** N = 39 participants (18 female) viewed 75 negative and 75 neutral IAPS images while EEG was recorded. After viewing the images, participants took a two-hour long nap which was followed by a memory assessment. Memory accuracy was measured using d’ = Z(hit rate) – Z(false alarm rate), where hit rate refers to the proportion of images seen during the memory assessment that are correctly identified as being previously seen, false alarm rate refers to the proportion of images seen during the memory assessment that are incorrectly identified as being previously seen, and Z() is the inverse cumulative distribution function of the standard normal distribution function.

**Results:** The early (300 – 1000 ms) and late (1000 – 1500 ms) LPP mediated enhanced discrimination accuracy for emotional compared to neural stimuli (d’) (*ps* < 0.001). The association between the late LPP and d’ was moderated by sleep such that the association was stronger when participants spent proportionately more time in N3 and REM (*p* = 0.02). The differences in reactivity between emotional and neutral images for both the early and late LPP were attenuated in PTSD+ individuals vs. controls (*ps* < 0.001). Despite mediation results showing greater d’ for emotional compared to neutral stimuli, women showed overall worse memory accuracy for negative compared to neutral stimuli (*p* < 0.001) whereas men showed no difference (*p* = 0.64).

**Conclusions:** N3 and REM sleep play a critical role for memory of stimuli that produce large and sustained neural responses. PTSD is marked by a diminished ability to distinguish between negative and neutral information. More research is critical to understand sex effects on emotional memory.

## Introduction

A rich body of laboratory research has established that information that stirs up emotions is better recalled than information that is experienced as emotionally neutral (Brown & Kulik, 1977; for reviews see Hall et al., 2021; Hamann, 2001; Kensinger & Corkin, 2003; LaBar & Cabeza, 2006; Mather & Nesmith, 2008; Phelps, 2006; Weymar et al., 2009; Yonelinas & Ritchey, 2015). While this emotional memory bias is thought to confer a survival benefit, it also has implications for psychiatric disorders, which is no better highlighted than in the case of posttraumatic stress disorder (as reviewed in Durand et al., 2019). The symptoms of posttraumatic stress disorder are anchored to the memory of a highly stressful life event. Individuals with PTSD are plagued with persistent, distressing memories of the traumatic event (American Psychiatric Association, 2013) that manifest during both wake and sleep (Richards et al., 2025), and they also typically experience fear states (physiological distress and psychic anxiety) when triggered by external reminders of their lived trauma (Glad et al., 2017; Glad et al., 2016; Glad et al., 2023). Understanding the processes involved in the encoding and consolidation of emotionally salient events in individuals with PTSD is therefore of critical importance.

Sleep has been demonstrated to play an important role in memory processing and also appears to have an important function in terms of emotion processing, PTSD development and PTSD recovery. For instance, REM sleep has been proposed to be important in emotional information processing (for a review see Walker & van der Helm, 2009); and disruptions in REM sleep after trauma have been associated with development of PTSD (Mellman & Hipolito, 2006; Mellman et al., 2007). Implicit fear memory studies have highlighted that REM sleep is important for the occurrence and/or consolidation of extinction learning, which is the new learning that competes with associations between neutral stimuli and fearful responses after classical conditioning of fear (Menz et al., 2016; Richards et al., 2022). Extinction learning and its consolidation are thought to be some of the core processes that underlie recovery from PTSD (Smith et al., 2017), and that permit individuals to consolidate declarative memories of traumatic occurrences without the intense emotions and narrative incoherence they initially elicited. In addition to REM sleep, NREM sleep and several of its features have a well-established role in memory consolidation **(Crowley et al., 2024)**. Emerging research also indicates that NREM sleep, especially N3 (deep NREM) sleep, has a function in emotion processing and emotional memory consolidation in particular (Yuksel et al., 2025). Converging lines of evidence thus suggest that disturbed sleep after trauma exposure plays an important contributing factor in maladaptive memory processing and consolidation.

While accumulating evidence supports a role for sleep in emotional learning, the pathway from wake encoding to preferential sleep consolidation of emotional information remains opaque. Presumably, however, emotionally salient information is encoded in a manner unique from neutral information. Indeed, evidence for this includes (event related potential) ERP research demonstrating that emotional stimuli generate a larger neural response than do neutral stimuli (Hajcak & Foti, 2020). More specifically, the late positive potential of the ERP, a deflection in the cortical EEG that emerges approximately 300 milliseconds post stimulus onset, is increased in response to emotional stimuli relative to neutral stimuli (for a review see Hajcak & Foti, 2020). Furthermore, it has been shown that the magnitude of the LPP is correlated with subsequent stimulus memory (as reviewd in Fields, 2023). For example, Dolcos & Cabeza (2002) found that the ERP was more positive following emotional compared to neutral stimuli and that the ERPs were more positive for remembered compared to forgotten stimuli. While the LPP has typically been measured in the window between 300 and 1000ms post stimulus onset, there is clear evidence that it is more protracted (Cuthbert et al., 2000). Research to date, however, has not demonstrated how this initial encoding difference translates into a memory difference and how sleep might play a role in transferring information that is preferentially tagged at encoding for memory consolidation. Furthermore, this has not been examined in trauma-exposed individuals with and without PTSD, in whom understanding differences in emotional memory processes might be particularly illuminating.

The current study examines this question in a sample of men and women with a history of psychological trauma with and without PTSD. Using a laboratory protocol involving an image viewing followed by a nap, the current study examined whether features of sleep moderate the relationship between the ERP response at encoding and subsequent memory. More specifically, we tested the hypothesis that the proportion of N3 and REM sleep in a nap influences the degree to which the emotional-neutral LPP differential predicts an emotional memory benefit for memory accuracy (See Figure 1a for model depiction). Supplementing this primary question, and with the objective of understanding the role of PTSD status and biological sex on sleep-dependent processing of emotional information, we expanded our framework to a larger model that factors these two characteristics. It is well established that women are at higher risk of PTSD after trauma than men, even when controlling for type of trauma. Sex differences in neural responses to emotional stimuli and differences in the role of sleep in the processing of those responses has not been adequately examined. The overall model proposes that negative (emotional) stimuli will be recalled better than neutral stimuli, and that the magnitude of the LPP mediates that effect and is moderated by sleep (High N3 and REM vs. Low N3 and REM). We then further explored whether PTSD status (positive vs. negative) and biological sex (male vs. female) moderated the relationship between stimulus type and neural response (See Figure 1b for the overall model). While we predicted that females would have a stronger LPP to negative relative to neutral images, our predictions with respect to PTSD were exploratory. Individuals with PTSD have been demonstrated to have a higher reactivity to negative images but also a higher reactivity to neutral images, which would blunt relative reactivity to neutral images.

**Figure 1.**
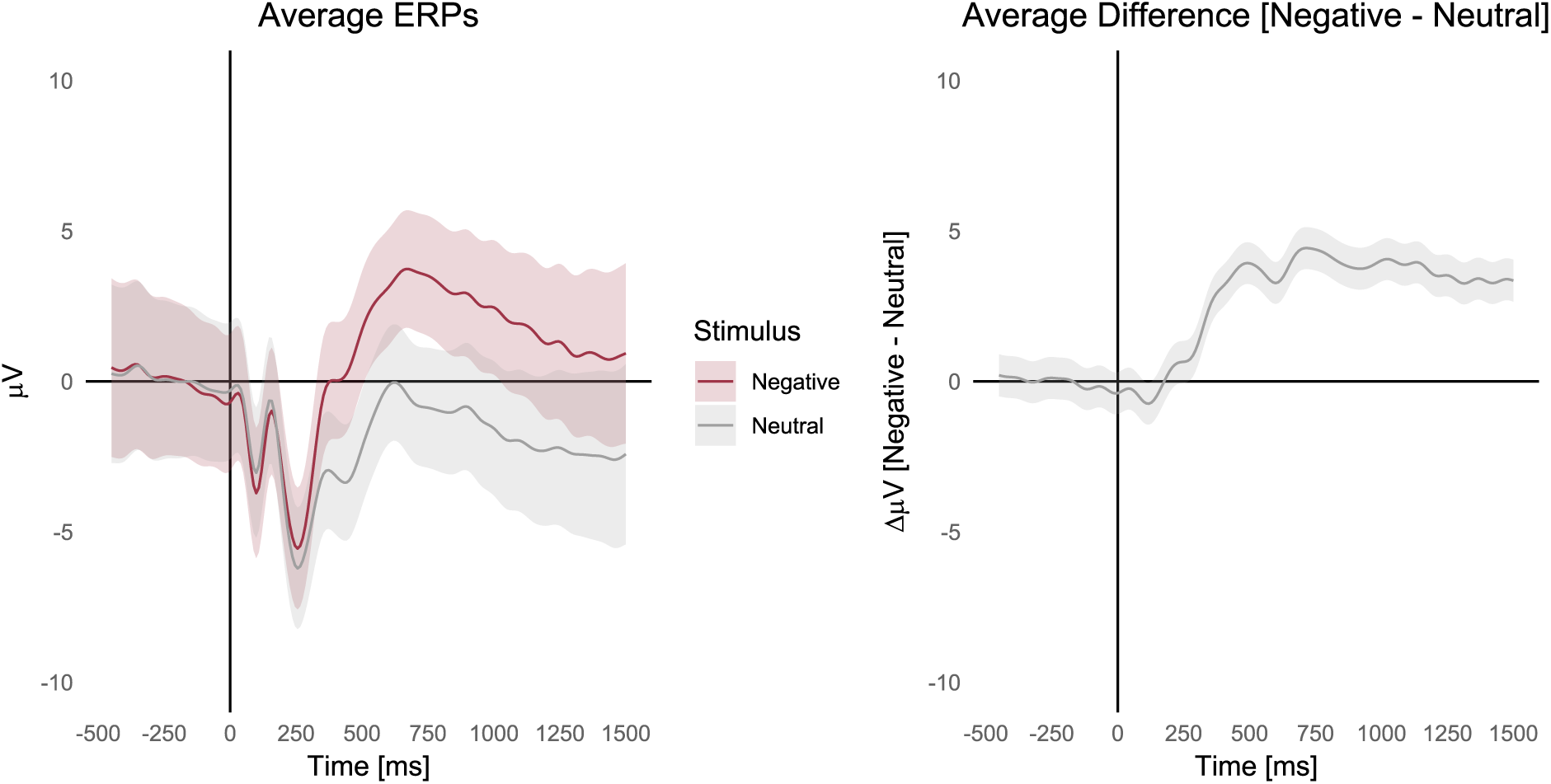
Generalized additive mixed effects models of the grand average ERP waveforms (left) and condition differences curves (right). Confidence regions depict 95% simultaneous confidence intervals across the entire curve. Averages correspond to averages across the four electrode sites (Cz, Pz, CP1, and CP2).

## Methods and Materials

### Participants

46 participants with a history of criterion A trauma exposure and between the ages of 18 and 50 were recruited from the San Francisco VA Medical center. Of these 46, N = 39 (18 female) were analyzed (n = 4 were excluded due to poor EEG signals and n = 3 were removed as outliers based on visual inspection of the data) (Table 1). We note that the results were not qualitatively impacted by the exclusion of the three outliers.

**Table 1.** Participant demographics and clinical features.

| Characteristic | N = 39 |
| --- | --- |
| Age, mean (SD) | 32.8 (5.9) |
| Biological sex, n (%) |  |
| Female | 18 (46.2) |
| Male | 21 (53.8) |
| Race, n (%) |  |
| White | 13 (33.0) |
| Asian | 10 (25.6) |
| Black or African American | 3 (7.7) |
| American Indian/Alaska Native | 1 (2.6) |
| Other | 10 (25.6) |
| Unknown | 2 (5.0) |
| Ethnicity, n (%) |  |
| Hispanic or Latino | 8 (20.5) |
| Not Hispanic or Latino | 28 (71.8) |
| Unknown | 3 (7.7) |
| CAPS-5 score, mean (SD) | 15.0 (11.0) |
| PTSD status, n (%) |  |
| PTSD+ | 12 (30.8) |
| PTSD- | 27 (69.2) |

### Experimental design

Participants viewed 75 negative and 75 neutral IAPS images (Lang et al., 1997) while electroencephalography (EEG) was recorded (Natraj et al., 2023). Each image was presented for 1500ms, followed by a fixation cross of variable duration (see Natraj et al., 2023). Following the image viewing session, participants had the opportunity to take a 120-minute nap between 1:30 and 3:30 PM, during which polysomnographic data were recorded from which sleep staging was scored (Natraj et al., 2023). Participants’ memories for the images were evaluated after the nap by showing them each of the 150 images that they had viewed and an additional 60 new images. The combined set was pseudo-randomly re-ordered. After each image presentation participants responded as to whether the image was previously seen (“old”) or not (“new”).

### Electroencephalography signal analyses

EEG was recorded at 1024 Hz with electrodes at four sites (Cz, Pz, CP1, and CP2). The four sites were chosen based on previous research indicating where the late positive potential is most pronounced (Hajcak & Foti, 2020). Raw EEG data were first band-pass filtered to 0.1 to 20Hz. Data were then re-referenced to the average of the mastoids and epochs extracted around stimulus presentations from −500ms to 1500ms. Blinks and eye artifacts were then corrected in the data using the recursive least squares algorithm (RLS) with the EOG leads as noise sources (a forgetting factor of 0.9999, and filter order of 3 (Natraj et al., 2018, He et al, 2004, Gomez-Herrero 2007, Hayes 2009). Epoched EEG data were then baselined to the first 500ms. In addition to the artifact correction, bad trials at each channel were identified using a threshold procedure. Specifically, a trial was deemed to be bad if at any point the ERP exceeded 5 S.D. of the mean ERP value across all trials for that specific channel. The channel-specific identified bad trials were then removed from further analyses. EEG data were down sampled to 40 Hz prior to entry in the generalized additive mixed models described below.

### Electroencephalography Sleep Data Acquisition

Polysomnographic methods are described in detail in (Natraj et al., 2023). Briefly, polysomnographic EEG data were collected at 400 Hz at locations conforming to the standard 10-20 electrode configuration with electrodes over frontal (F3, F4), central (C3, C4), and occipital (O1, O2) sites. Also collected were 3 bipolar chin electromyograms, horizontal electro-oculography, and electrocardiogram. EEG and electro-oculography signals were referenced to the contralateral mastoid electrodes and then re-referenced to a linked mastoid reference. Signals were then filtered at 0.3 to 35 Hz and sleep stages (N1, N2, N3, or rapid eye movement) were scored in 30-second epochs according to the American Academy of Sleep Medicine criteria using the PRANA Software Suite, version 10 (PhiTools).

## Statistical Methods

All analyses were conducted in R version 4.3.3 (R Core Team, 2024).

### Evaluating the late positive potential (LPP)

Differences in the ERP 300 – 1000 ms following stimulus onset (time window chosen *a priori*) were assessed using linear mixed effects models using the lme4 R package (Bates et al., 2015). Analyses were conducted at the trial level while including random intercepts for both participants and stimuli (crossed random effects). The average ERP across the time window of interest within each electrode site on each trial was used as the dependent variable with the independent variable being stimulus type (negative vs neutral) while adjusting for sex, PTSD status, age, and electrode site. Sex and PTSD differences in neural reactivity to the stimuli was assessed by including interactions between the variable of interest and stimulus type. Females and PTSD-individuals were used as the reference category in all models. Significance was declared at the α = 0.05 level and parameter estimates along with their standard errors (in parentheses), test statistics, and p-values are reported for all models.

While we chose the 300 – 1000 ms window *a priori* based on a predominance of research focused on this window, negative vs. neutral differences in the LPP have been observed to last beyond that window (Cuthbert et al., 2000), and these might have implications for encoding and memory. We were therefore interested in examining condition differences for the duration of image viewing and we used a data-driven approach to investigate this. To do so, we fit generalized additive mixed effects models (GAMMs) (Wood, 2017) to the ERP at the trial level to model the entire ERP waveform from −500 ms before to 1500 ms after stimulus onset. See the Supplemental Methods for details of the GAMM specification. This allowed us to evaluate effects at two separate time windows, 300 – 1000 ms (the early LPP) and 1000 – 1500 ms (the late LPP).

### Memory assessment

Memory accuracy as measured by d’ = *Z*(Hit Rate) – *Z*(False Alarm Rate) was of primary interest. Here, hit rate corresponds to the proportion of trials on which participants reported an image as being seen during encoding among images that were in the encoding set. False alarm rate corresponds to the proportion of trials on which participants reported an image as being seen during encoding among images that were not in the encoding set. *Z*() corresponds to the inverse cumulative distribution function of the standard normal distribution. Linear mixed effects models were used to test for stimulus type differences in d’, Hit Rate, and False Alarm Rate with random intercepts given for participants. Sex and PTSD differences were assessed by including sex * stimulus type and PTSD status * stimulus type interactions in the models and all models adjusted for age.

### Sleep composite score

A sleep composite score (Richards et al., 2022) was computed by subjecting the proportion of time spent awake and in N1, N2, N3, and REM sleep to factor analysis for compositional data using the *pfa()* function in the robCompositions R package (Templ et al., 2011). The first latent factor explained 31% of the variance in the observed data and took on higher values when participants spent proportionately more time in N3 and REM sleep and less time awake and in N1 (Table 2). This factor, distinguishing between proportionally high N3 and REM sleep (referred to as high N3/REM here) and low N3 and REM sleep (referred to as low N3/REM here) was correlated with subjective sleep quality (Richards et al., 2022), and contrasts sleep rich in features of the sleep period considered important for emotional memory (N3 and REM) from N1 and wake. This factor was used to generate each subject’s *sleep composite score* and was standardized to have a mean of 0 and SD of 1 prior to entry in all models.

**Table 2.**
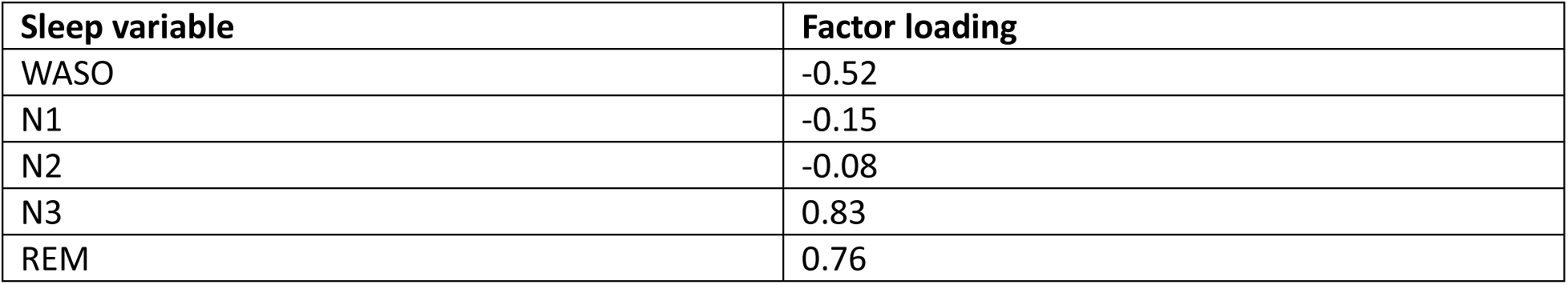
Compositional factor analysis loadings for the sleep composite score. Participants who spent proportionately more time in N3 and REM and less time in WASO and N1 have greater values of the sleep composite score. WASO = wake after sleep onset, REM = rapid eye movement.

### Multilevel mediation analyses

Multilevel mediation analyses were performed following the methods of Zhang et al. (2009) to test the hypothesis that differences in memory accuracy (d’) for negative compared to neutral stimuli were mediated by the LPP. Specifically, linear mixed effects models were used to model the LPP at the trial level in the 300 – 1000 ms window following stimulus onset (mediation analysis path *a*) while adjusting for channel. Generalized linear mixed effects models (GLMMs) were then used to test for stimulus type differences in d’ at the trial level (path *c’*) while adjusting for the average subject-mean-centered LPP in the 300 – 1000 ms across all 4 channels (path *b*) (Zhang et al., 2009). Specifically, binomial GLMMs with a probit link function modeled whether the participant correctly identified that the stimulus was seen before (a “hit”) as the dependent variable while including an offset term of the Z-transformed false alarm rate for the stimulus type (negative or neutral). Inclusion of this offset term leads to a model parameterization where the dependent variable is d’. Random intercepts were given for both subjects and stimuli (crossed random effects). These two models were fit 10,000 times by parametric bootstrap using the *bootMer*() function in the lme4 (Bates et al., 2015) package from which direct (path *c’*), indirect (path *a* * *b*), and total (c’ + a * b) effects of stimulus type on d’ were computed. All models adjusted for sex, PTSD status, and age.

Moderated mediation analyses were performed to test the hypotheses that the direct, indirect, and total effects of stimulus type on d’ were moderated by the sleep composite score, sex, and PTSD status. The moderated mediation analyses were accomplished by including interactions between these variables and the appropriate paths along the mediation model (paths *a*, *b*, and *c’*). Models were fit 10,000 times using the parametric bootstrap and differences in the bootstrapped conditional indirect, direct, and total effects were computed for sex, PTSD status, and the sleep composite score.

## Results

### The late positive potential (LPP)

Results of the GAMM demonstrated that the stimulus differences in the ERP began 290 ms post stimulus onset and were sustained for the duration of stimulus presentation, with the ERP being more positive for negative compared to neutral stimuli (the *late positive potentia*ļ LPP) (*p* < 0.001) (Figure 1). While the ERP amplitudes themselves did differ across electrode sites (*p* < 0.001), the condition difference between negative and neutral stimuli did not (*p* = 0.18, Supplementary Figure 1) therefore we present results as the averages over the four electrodes.

Consistent with the results above, the amplitude of the early LPP was larger following negative compared to neutral stimuli in the 300 – 1000 ms window post-stimulus onset (*b* = 3.61 (0.62), t(147.1) = 5.81, *p* < 0.001). The late LPP (1000 – 1500 ms post stimulus) was also larger following negative compared to neutral stimuli (*b* = 3.74 (0.51), *t*(147.58) = 7.41, *p* < 0.001).

### The LPP predicts memory accuracy and the association between the late LPP and d’ is moderated by properties of sleep

We next sought to test the hypotheses that the early and late LPP were associated with memory accuracy (d’). Here, we found that both early (*b =* 0.008 (0.002), *z* = 3.55, *p* < 0.001) and late LPP (*b =* 0.005 (0.002), *z* = 3.08, *p* < 0.001) were predictive of d’.

While the early LPP was predictive of d’, this association was not moderated by the sleep composite score (*b* = 0.002 (0.002), *z* = 0.98, *p* = 0.33). In contrast, the association between the late LPP and d’ was significantly moderated by the sleep composite score (*b* = 0.004 (0.002), *z* = 2.34, *p* = 0.02) (Figure 2). Specifically, there was no association between the late LPP and d’ in participants who spent proportionately less time in N3 and REM sleep (*b* = 0.001 (0.002), *z* = 0.32, p = 0.75), but there was in participants who spent proportionately more time in these sleep stages (*b* = 0.01 (0.002), *z* = 3.72, *p* < 0.001). Put another way, memory accuracy increased with neural reactivity during the late LPP but only when participants spent proportionately more time in N3 and REM sleep. Post-hoc analyses showed that this moderation effect was largely driven by N3 sleep (*p* = 0.02), and not by REM (*p* = 0.23).

**Figure 2.**
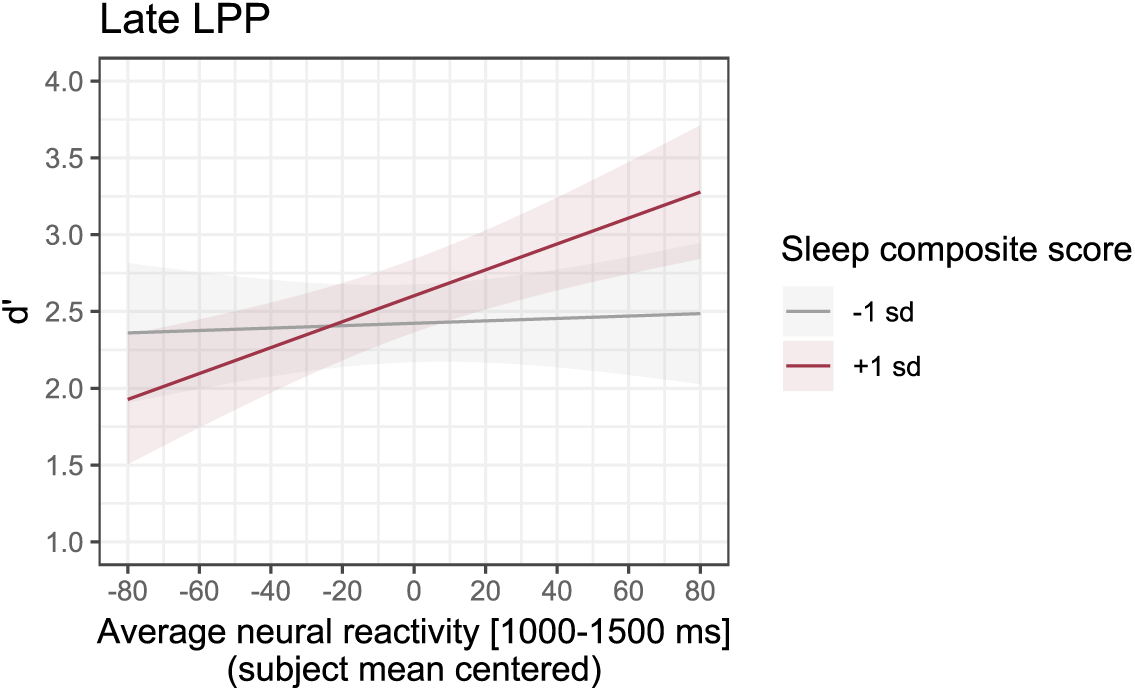
The association between the late LPP and memory accuracy. (d’) was moderated by the standardized sleep composite score (p = 0.02) (mean – 1SD and mean + 1SD are shown in grey and red, respectively), controlling for sex and PTSD. Greater values of the sleep composite score indicate proportionately more time spent in N3 and REM sleep during the nap.

As an exploratory analysis as to exactly when this interaction emerged in the LPP, we computed the average LPP every 25 ms using a sliding window with a width of 50 ms. The Z-statistics for the interaction along this sliding window were subjected to threshold-free cluster enhancement (Smith & Nichols, 2009) and p-values were adjusted for multiple comparisons across the entire LPP. Results suggested that the interaction effect was significant in a window from 950 to 1350 ms post-stimulus onset (Supplementary Figure 2), which matches well with the 1000 – 1500 ms window used for the late LPP above.

### An emotional memory benefit is mediated by both the early and late LPP

While the results above suggest a possible mediation of differences in d’ due to differences in the LPP they do not formally test this hypothesis. To do so, we performed formal mediation analyses to test for stimulus type differences in d’ that were mediated by the early and late LPP (Figure 3). The early LPP was more positive following negative compared to neutral stimuli (*b* = 3.61 (0.62), t(147.1) = 5.81, *p* < 0.001) and was positively predictive of d’ (*b* = 0.008 (0.002), z = 3.55, *p* < 0.001) resulting in a significant positive indirect effect of stimulus type on d’ (*indirect effect* = 0.029, 95% bootstrapped CI: [0.012, 0.049], *p* < 0.001). That said, the direct effect of stimulus type on d’ was *negative* such that, adjusting for the early LPP, memory accuracy was worse for negative compared to neutral stimuli (*b* = −0.22 (0.11), z = −2.10, *p* = 0.04). Consequently, the total effect of stimulus type on d’ was negative, indicating that d’ was overall lower for negative compared to neutral stimuli (*total effect* = −0.20, 95% bootstrapped CI: [−0.40, −0.003], *p* = 0.048).

**Figure 3.**
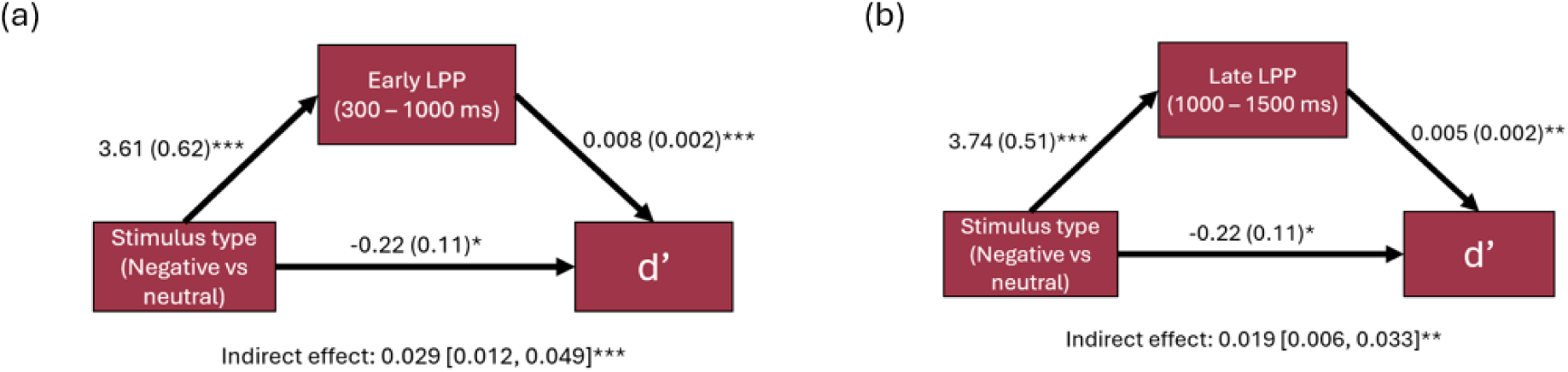
Mediation models of the (a) early and (b) late LPP. Parameter estimates with their standard errors in parentheses are shown. Models adjusting for sex, PTSD status, and age. An emotional memory benefit was mediated by both the early and late LPP. * p < 0.05, ** p < 0.01, *** p < 0.001.

Exploratory analyses of the late LPP resulted in similar findings (Figure 3b). That is, the late LPP was more positive following negative compared to neutral stimuli (*b* = 3.74 (0.51), *t*(147.58) = 7.41, *p* < 0.001), the late LPP was positively predictive of d’ (*b* = 0.005 (0.002), z = 3.08, *p* = 0.002), and there was a significant indirect effect of the late LPP on d’ (*indirect effect* = 0.019, 95% bootstrapped CI: [0.006, 0.033], *p* = 0.002). Again, the total effect was negative such that overall, d’ was lower for negative compared to neutral stimuli (*total effect* = −0.20, 95% bootstrapped CI: [−0.40, −0.01], *p* = 0.046).

### Properties of sleep moderate the indirect effect of stimulus type on d’ as mediated by the late LPP

Given that the association between the late LPP and d’ was found to be moderated by the sleep composite score, we wished to perform a moderated mediation analysis to see if the emotional memory benefit was moderated by properties of sleep. Results of these analyses revealed that there were no significant differences in the indirect effects of the early LPP (*high N3/REM – low N3/REM indirect effect* = 0.014 95% bootstrapped CI [−0.016, 0.047], *p* = 0.37) (Figure 4a), but there were differences in the indirect effects of the late LPP (*high N3/REM – low N3/REM indirect effect* = 0.025 95% bootstrapped CI [0.001, 0.051], *p* = 0.039) (Figure 4b). Specifically, there was no indirect effect on d’ in participants who spent less time in N3/REM (*indirect effect =* 0.006, 95% bootstrapped CI [−0.012, 0.024], *p* = 0.53) but there was an indirect effect in those who spent proportionately more time in N3/REM (*indirect effect =* 0.031, 95% bootstrapped CI [0.013, 0.051], *p* < 0.001). Put another way, an emotional memory benefit as mediated by the late LPP emerged only when participants spend more time in N3/REM sleep.

**Figure 4.**
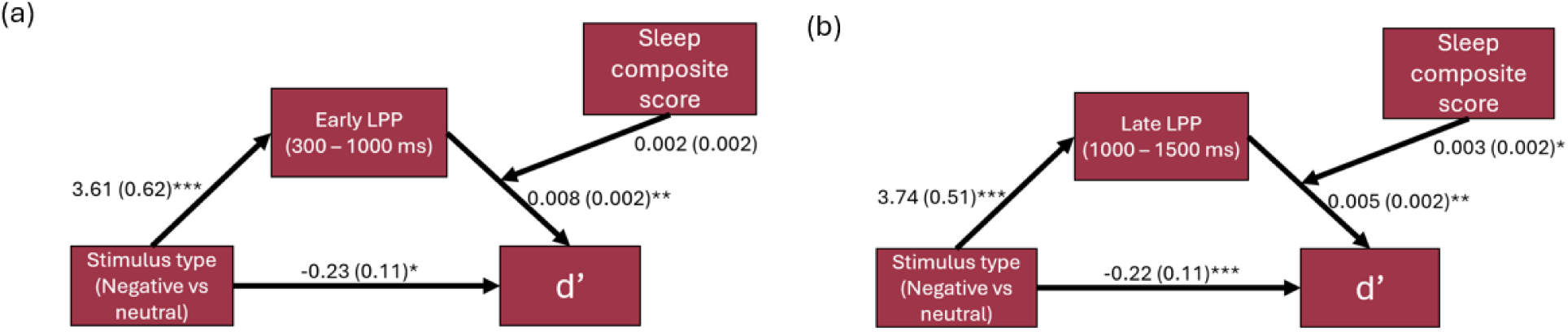
Moderated mediation models of the (a) early and (b) late LPP. Parameter estimates with their standard errors in parentheses are shown. The indirect effect of the late LPP was moderated by the sleep composite score. See the text for the conditional indirect effects. Models adjusted for sex, PTSD, and age. An emotional memory benefit was mediated by both the early and late LPP. * p < 0.05, ** p < 0.01, *** p < 0.001.

### Both the early and late LPP were moderated by PTSD status but not by sex

We next sought to test for PTSD differences in the LPP. Here, PTSD status was found to moderate both the early (*b* = −1.20 (0.33), *t*(21760) = −3.66, *p* < 0.001) and late LPP (*b* = −1.73 (0.44), *t*(21760) = −3.89, *p* < 0.001), that is, there was less of a difference between the negative and the neutral stimulus responses in PTSD+ individuals (Figure 5). Sex was not found to significantly moderate the early (*b* = −0.24 (0.3), *t*(21760) = −0.82, *p* = 0.41) or the late LPP (*b* = 0.41 (0.41), *t*(21764) = 1.0, *p* = 0.32) (Figure 6).

**Figure 5.**
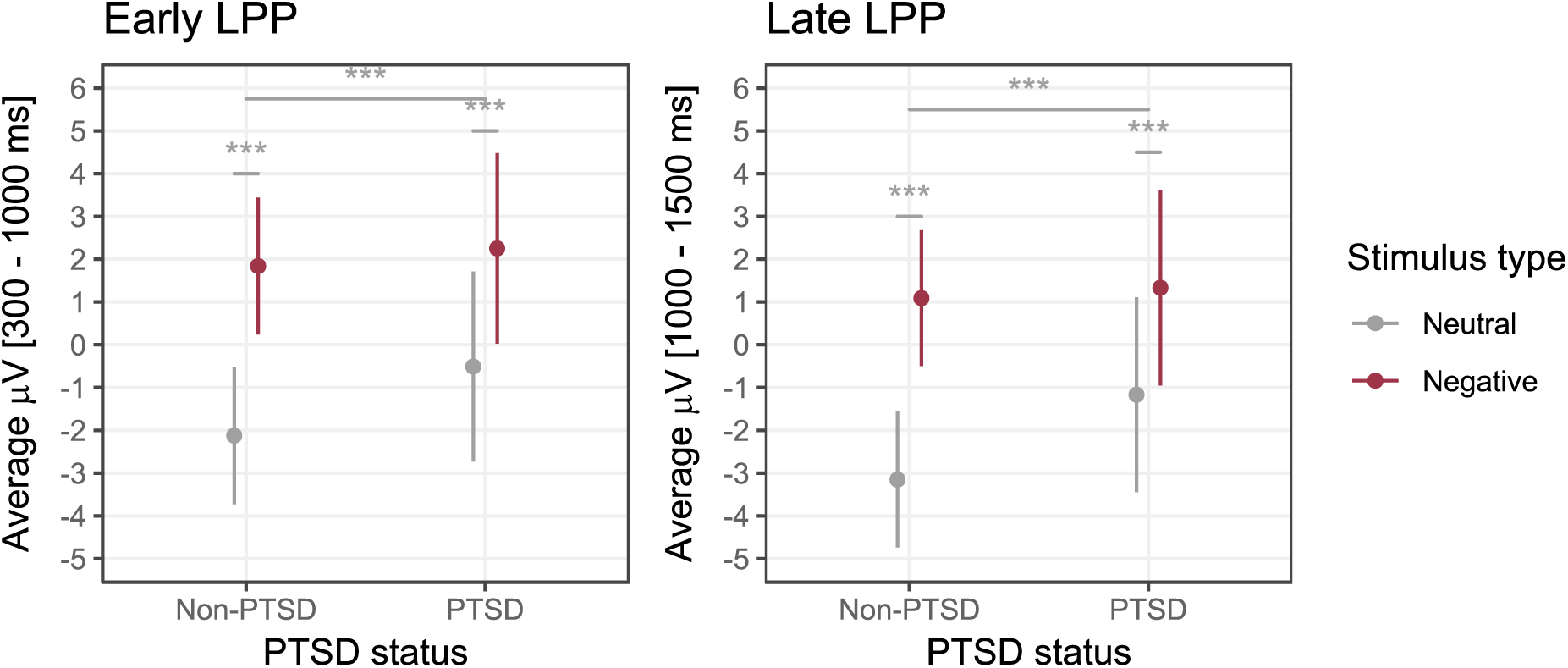
Late positive potential (LPP) by PTSD status. The early LPP (300 – 1000 ms) is shown on the left and the late LPP (1000 – 1500) is shown on the right. Models adjusted for sex and age. * p < 0.05, ** p < 0.01, *** p < 0.001, NS = Not Significant. The top bar in each panel corresponds to the significance level of the PTSD status * stimulus type interaction.

**Figure 6.**
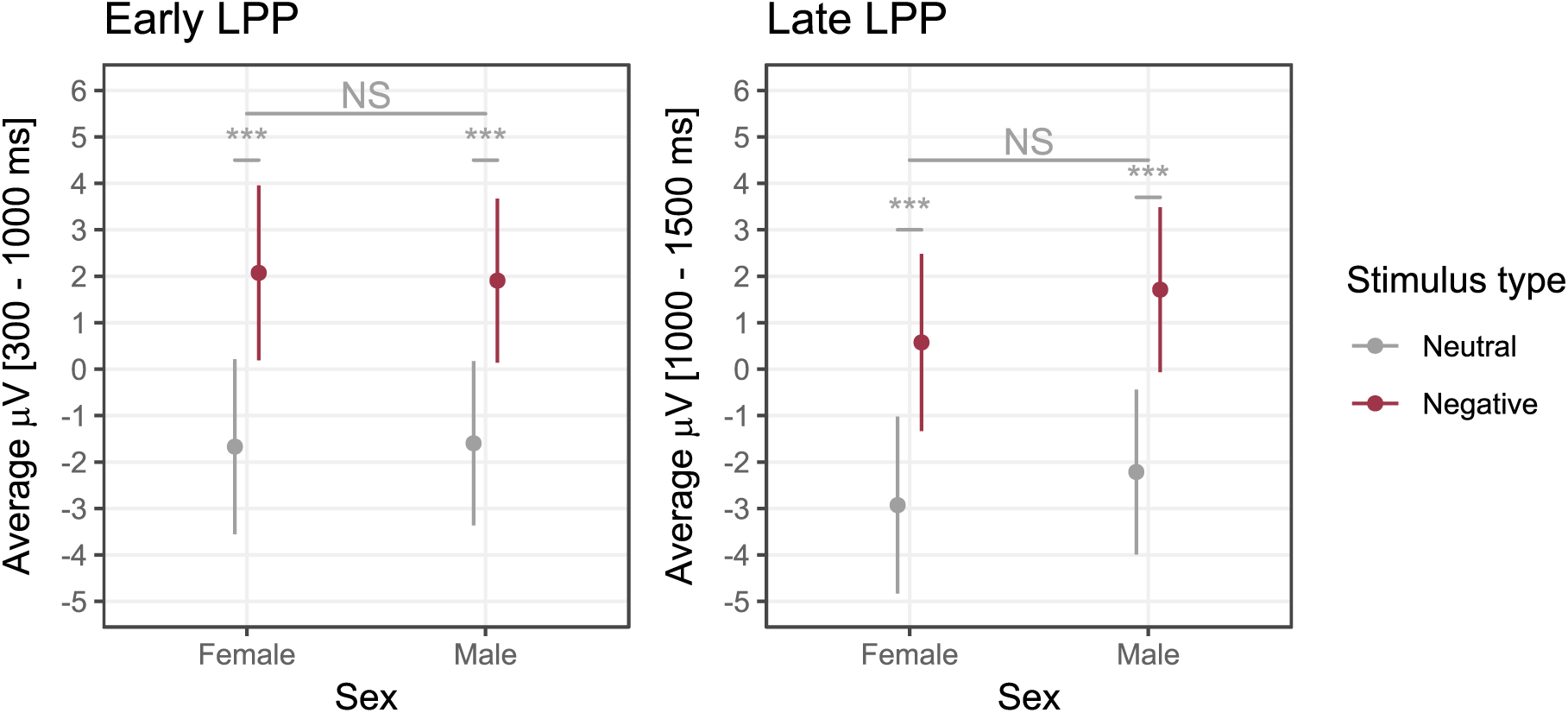
The late positive potential (LPP) by sex. The early LPP (300 – 1000 ms) is shown on the left and the late LPP (1000 – 1500) is shown on the right. Models adjusted for PTSD status and age. * p < 0.05, ** p < 0.01, *** p < 0.001, NS = Not Significant. The top bar in each panel corresponds to the significance level of the sex * stimulus type interaction.

### Memory accuracy was moderated by sex but not by PTSD status

Further exploratory analyses investigated sex and PTSD differences in memory. Sex was found to moderate d’ (*p* = 0.005) with women showing *lower* d’ for negative compared to neutral stimuli ( = −0.49 (0.13), *t*(41) = −3.66, *p* < 0.001) and men showing no difference (*b* = 0.058 (0.12), *t*(41) = 0.47, *p* = 0.64) (Figure 7). Further, sex was found to moderate hit rate (*p* = 0.001) but not false alarm rate (*p* = 0.61). Here, women showed no significant difference in hit rate between negative and neutral stimuli (*b* = −0.013 (0.02), *t*(41) = −0.63, *p* = 0.53) despite an increased false alarm rate for negative compared to neutral stimuli (*b* = 0.065 (0.02), *t*(41) = 3.23, *p* = 0.002). That is, the lower d’ for emotional compared to neutral stimuli in women was due to an increased false alarm rate for negative relative to neutral images which was not followed by an increase in hit rate for negative compared to neutral stimuli. In contrast, men showed a significantly greater hit rate (*b* = 0.084 (0.019), *t*(41) = 4.49, *p* = 0.001) and false alarm rate (*b* = 0.079 (0.019), *t*(41) = 4.21, *p* = 0.001) for negative compared to neutral stimuli. That said, there were no significant differences between men and women when directly contrasting negative image d’, hit rate, and false alarm rate (*ps* > 0.05). There were similarly no significant sex differences when directly contrasting d’, hit rate, and false alarm rate for neutral images (*ps* > 0.05).

**Figure 7.**
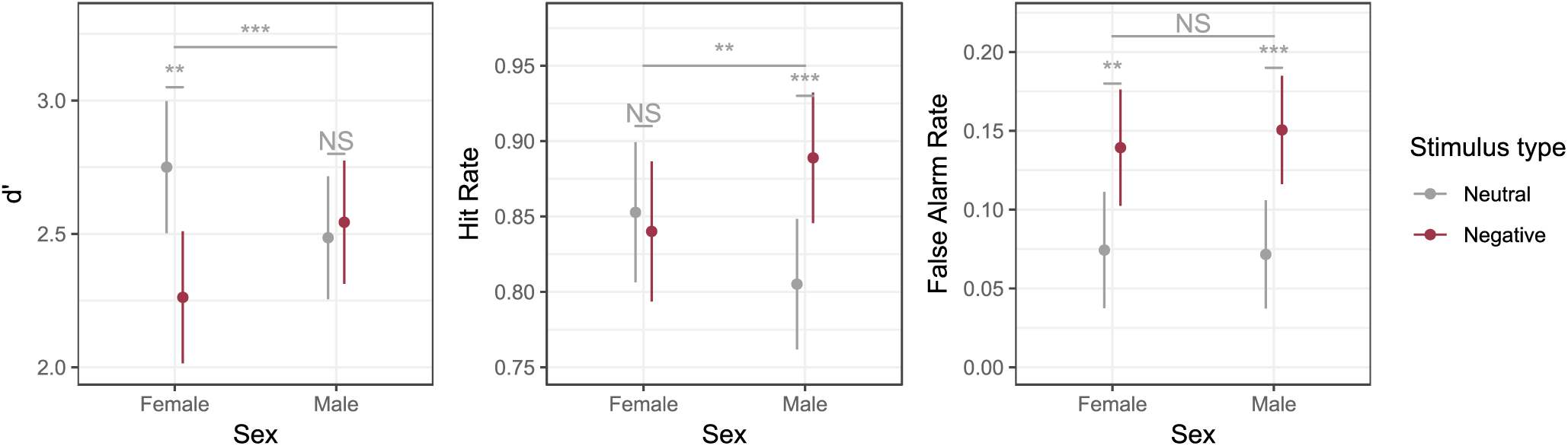
Memory performance by sex. The top bar in each panel corresponds to the significance level of the sex * stimulus type interaction. Models adjusted for PTSD status and age. * p < 0.05, ** p < 0.01, *** p < 0.001, NS = Not Significant.

PTSD status was not found to moderate d’ (*p* = 0.55), hit rate (*p* = 0.23), or false alarm rate (*p* = 0.32) while adjusting for the sex * stimulus type interaction in each model (Figure 8).

**Figure 8.**
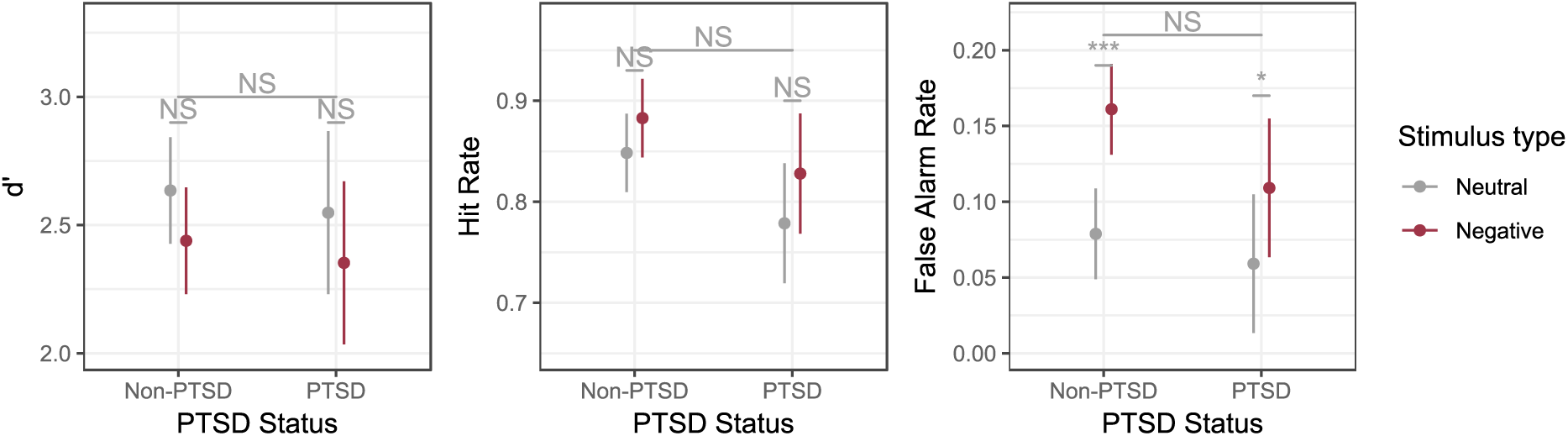
Memory performance by PTSD status. The top bar in each panel corresponds to the significance level of the PTSD * stimulus type interaction. Models adjusted for sex and age. * p < 0.05, ** p < 0.01, *** p < 0.001, NS = Not Significant.

### Sex moderates the direct effect of stimulus type on d’ but does not impact the indirect effect as mediated by the late LPP

We next sought out to test the hypothesis that sex moderated the indirect, direct, and total effects of stimulus type on d’ as mediated by the early LPP. While sex was not found to significantly moderate the early LPP (*b* = −0.24 (0.3), *t*(21760) = −0.82, *p* = 0.41), sex did significantly impact the direct effect (*b* = 0.65 (0.10), *z* = 6.75, *p* < 0.001). That is, 1) there were no significant sex differences in the early LPP, 2) there was a significant indirect effect of stimulus type on d’ through the early LPP for *both* women (*indirect effect* = 0.028, 95% bootstrapped CI [0.011, 0.049], *p* < 0.001) and men (*indirect effect* = 0.026, 95% bootstrapped CI [0.010, 0.046], *p* < 0.001), and 3) this indirect effect did not differ between sexes (*Male - Female indirect effect* = −0.002, 95% bootstrapped CI [−0.007, 0.003], *p* = 0.42). That said, the direct effect did differ between sexes such that adjusting for the early LPP, women again evidenced significantly lower d’ for negative compared to neutral stimuli (*direct effect* = −0.56, 95% bootstrapped CI [−0.78, −0.33] *p* < 0.001) whereas men showed no difference (*direct effect* = 0.08, 95% bootstrapped CI [−0.14, 0.30], 95%, *p* = 0.50). Regarding the total effect, women had overall lower d’ for negative compared to neutral stimuli (*total effect* = −0.53, 95% bootstrapped CI [−0.75, −0.31], *p* < 0.001) whereas men showed no significant difference (*total effect* = 0.10, 95% bootstrapped CI [−0.12, 0.33], *p* = 0.36).

Similar results were observed for the late LPP (See Supplementary Results).

### PTSD status moderates the indirect effect of stimulus type as mediated by the LPP

Next, we tested the hypothesis that PTSD status moderated the degree to which differences in d’ were mediated by the LPP by including PTSD interactions with the indirect and direct effects. Models were adjusted for the sex * direct effect interaction that was found above. Results of these analyses showed the early LPP differed by PTSD status with the difference between negative and neutral images in the early LPP being lower for PTSD+ compared to PTSD-individuals (*b* = −1.20 (0.33), *t*(2176) = −3.66, *p* < 0.001). In contrast, the direct effect did not differ between PTSD+ and PTSD-individuals (*b* = 0.17 (0.10), *z* = 1.59, *p* = 0.11) therefore this moderation effect was not included in subsequent mediation models. The early LPP was found to mediate greater d’ for negative compared to neutral images for both PTSD-(*indirect effect* = 0.029, 95% bootstrapped CI [0.010, 0.051], *p* = 0.002) and PTSD+ individuals (*indirect effect* = 0.020, 95% bootstrapped CI [0.01, 0.038], *p* = 0.002) however the magnitude of the indirect effect was lower for PTSD + individuals (PTSD+ - PTSD-indirect effect = −0.01, 95% bootstrapped CI [−0.017, −0.003], *p* = 0.002).

Similar results were found for the late LPP (See Supplementary Results).

### Overall moderated mediation model

The full model which incorporated the moderation effects of the sleep composite score, PTSD status, and sex is shown in Figure 9. This model summarizes all of the aforementioned results: there is an emotional memory benefit mediated by both the early LPP and late LPP amplitudes (*ps* < 0.05) and this benefit is blunted in PTSD+ individuals due to hyperarousal to neutral stimuli (*ps* < 0.001). For the late LPP, the emotional memory benefit is enhanced when participants spend more time in N3/REM sleep (*p* < 0.05). Lastly, women have overall worse memory accuracy for negative images (*ps* < 0.001) and this is not due to differences in either the early or late LPP.

**Figure 9.**
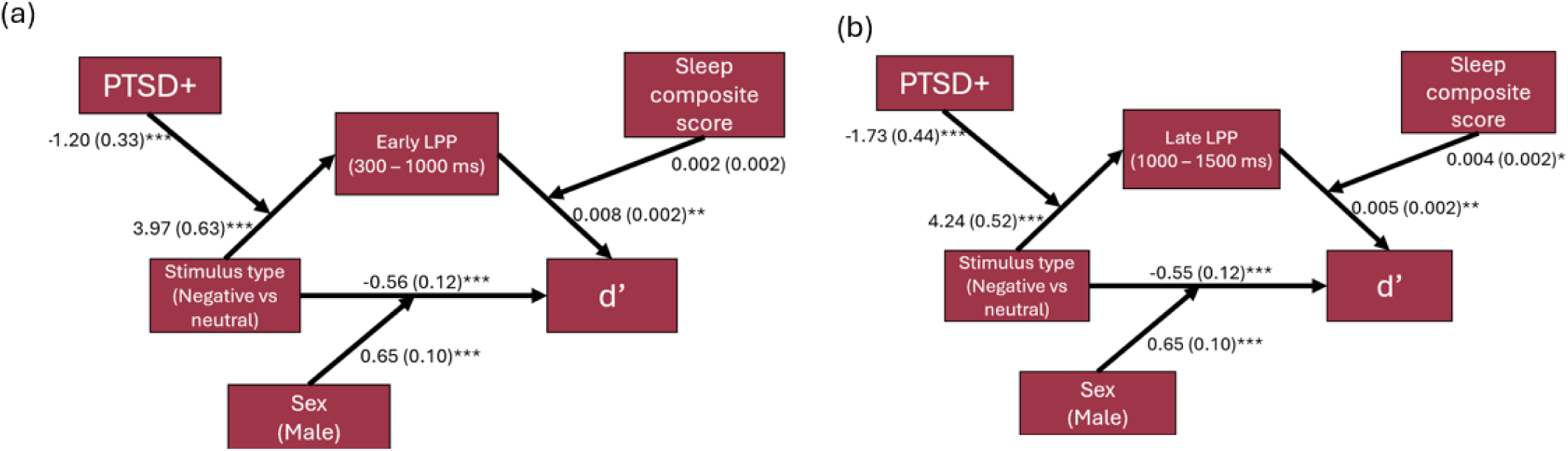
Moderated mediation models of the (a) early and (b) late LPP. Parameter estimates with their standard errors in parentheses are shown. An emotional memory benefit was mediated by both the early and late LPP. PTSD status moderated both the early and late LPP and sex moderated the direct effects of stimulus type on memory accuracy (d’). The sleep composite score moderated the association between the late LPP and d’. See the main text for the conditional indirect and total effects. * p < 0.05, ** p < 0.01, *** p < 0.001

## Discussion

Our analyses demonstrated that the late positive potential (LPP) mediates an emotional memory benefit and that, for the late LPP, only, this mediation is moderated by the proportion of time spent in N3 and REM sleep. That is, when participants spend proportionately more time in N3 and REM sleep there is a stronger association between the magnitude of the late LPP and memory accuracy. Sex differences in the total effect for emotional memory were observed with women showing worse memory accuracy for negative compared to neutral stimuli—driven by a greater false alarm rate for negative images but approximately equivalent hit rate between image types —whereas men showed no overall difference in d’. PTSD+ individuals evidenced a blunted difference in reactivity between negative and neutral stimuli resulting in a diminished emotional memory benefit in mediation analyses. These results inform the functional role of the LPP highlighting its importance in emotional memory, provide novel evidence as to the role that sleep plays in moderating this effect, and highlight sex and PTSD differences in emotional memory. Our results suggest that interventions targeting sleep architecture may be potential avenues for the treatment of PTSD-related memory impairment.

While the LPP has consistently been demonstrated to serve as a marker of emotional reactivity and regulation across the lifespan (Calentino et al., 2025; Dennis & Hajcak, 2009; Desatnik et al., 2017; Meynadasy et al., 2022), the functional role of the neural events underlying the LPP remains an open question. Hajcak & Foti (2020) argue that the LPP reflects stimulus significance, which they define as relevance to “[activate] appetitive or aversive motivational systems in the brain”. Critically, these authors argue that the LPP and the widely studied P300 may have a shared neurobiological mechanism, namely the locus coeruleus/norepinephrine system (Hajcak & Foti, 2020). Fields (2023) largely agrees with this view but also, borrowing from the P300 literature, argues that the LPP serves to encode significant stimuli into memory. That is, Fields (2023) points to the large P300 literature that has demonstrated that this component is important for memory consolidation. Our results are consistent with the hypothesis that the LPP serves to assist in the encoding of emotional stimuli into memory (Aston-Jones & Cohen, 2005; Fields, 2023).

Our findings that the N3/REM sleep composite variable moderates the association between the late LPP (1000 – 1500 ms) and memory accuracy are consistent with research demonstrating the importance these two sleep stages play in memory consolidation. For example, one recent meta-analysis of 125 studies showed that targeted sleep disruption of both N3 and REM sleep impairs memory encoding and consolidation, with the effects being larger for N3 compared REM sleep (Crowley et al., 2024). A similar meta-analysis of 185 studies showed that total sleep deprivation impairs memory both before and after learning (Newbury et al., 2021). Further studies have shown that time spent in N3 sleep positively correlates with performance in episodic memory tasks (Faghel-Soubeyrand et al., 2025; Leong et al., 2021; Lokhandwala & Spencer, 2021; Scullin, 2013). Our results expand on this literature by demonstrating that sleep rich in N3 and REM serves to moderate the conditions under which neural reactivity at encoding translates to enhanced memory performance. That is, stimuli that produce large and sustained neural responses that are then followed by proportionately higher amounts of N3 and REM sleep are remembered best; however, if there are low amounts of N3 and REM, sustained neural responses have no bearing on memory performance. Post-hoc analyses demonstrated that this result was largely driven by N3 sleep. Future studies with a full night of sleep may be more revealing of REM effects.

Critically, our finding that N3 and REM sleep moderated the association between the LPP and memory accuracy was specific to the late LPP (1000 – 1500 ms) and not the early LPP (300 – 1000 ms). That is, regardless of N3/REM, stimuli that provoke strong initial neuronal responses are preferentially encoded into memory; however, stimuli that provoke sustained neuronal responses are only consolidated if they are then followed by proportionately more N3/REM sleep. While speculative, it may be that stimuli that produce sustained neural activation engage a broader and more complex set of neural networks that effectively tags them for sleep-dependent consolidation.

The finding that, for the total effect, emotional memory accuracy for emotional images was worse than that for neutral images in women but not in men stands in contrast to a large literature showing that emotional content is remembered better than neutral information (Brown & Kulik, 1977; for reviews see Hall et al., 2021; Hamann, 2001; Kensinger & Corkin, 2003; LaBar & Cabeza, 2006; Mather & Nesmith, 2008; Phelps, 2006; Weymar et al., 2009; Yonelinas & Ritchey, 2015). Moreover, these findings stand in contrast to a growing literature demonstrating that women have *superior* memory for emotional content compared to men (Bloise & Johnson, 2007; Canli et al., 2002; Felmingham et al., 2012). For example, women under stress recalled emotional images better than men (Felmingham et al., 2012), however the same study showed that women had worse memory for emotional images compared to men under a control condition (Felmingham et al., 2012). Given that enhanced memory for emotional images in women is only present in highly stressful environments (Felmingham et al., 2012) it may be that our emotional stimuli were not sufficiently arousing to cause the expected sex differences in memory. Another possible, but untested, explanation is that female subjects engaged in greater avoidance of negative stimuli during viewing, that might have contributed to lower hit rates observed here. Future research would benefit from eye tracking to measure position and duration of gaze during stimulus viewing.

Lastly, our finding that there was a blunted difference in the ERPs between negative and neutral stimuli in PTSD+ compared to PTSD-individuals is consistent with literature demonstrating either a hyperarousal to neutral stimuli in PTSD+ samples (Brunetti et al., 2010; Hendler et al., 2003; Kimble et al., 2000; Zukerman et al., 2018) or blunted reactivity to negative affective stimuli in PTSD+ individuals (DiGangi et al., 2018; DiGangi et al., 2017; Felmingham et al., 2003; MacNamara et al., 2013; Roeckner et al., 2025). For example, an fMRI analysis by Brunetti and colleagues (2010) found that, relative to PTSD-individuals, PTSD+ individuals evidence heightened BOLD response in the amygdala in response to neutrally valenced IAPS images. Further, there were no significant differences in BOLD activity in the amygdala between negative and neutral stimuli in PTSD+ individuals. Felmingham et a. (2003) found overall blunted ERP responses to angry and neutral faces in PTSD+ participants and the ERPs did not significantly differ between angry and neutral faces in PTSD+ participants. Our findings, together with the existing literature, suggest a diminished ability to discriminate between negative and neutral stimuli in PTSD as suggested in Felmingham et al. (2003). Moreover, our mediation analyses demonstrate that this failure to discriminate between negative and neutral stimuli in PTSD translates to a diminished emotional memory benefit in PTSD.

Our study has important limitations. First, even though we found that proportionally higher N3 and REM sleep moderated the relationship between the LPP and memory accuracy, this was based on a daytime nap rather than typical overnight sleep. Post-hoc examination of N3 and REM contributions separately indicated that N3 sleep accounted for the sleep moderation effects. We note that participants only reported 10 minutes of REM sleep on average, and while our prior analyses showed some effects of daytime nap REM sleep on implicit learning processes (Richards et al., 2022), such short REM sleep durations may have made it difficult to detect independent contributions of REM sleep in this moderation analysis. Future research investigating the relationships observed here using a full night of sleep with sleep stage durations representing those of a typical overnight sleep period is warranted. This is all the more important because PTSD sleep is well established to be deficient in N3 sleep; examining these relationships using overnight sleep would therefore further shed light on mechanisms explaining maladaptive memory consolidation in PTSD.

Another limitation of our analysis is that despite finding significant mediation effects for an emotional memory benefit, the total effect of the overall mediation model was non-significant. That said, it has been shown that under certain circumstances tests for mediated effects can have substantially higher power than tests for total effects (Kenny & Judd, 2014) which may be why we observed the significant mediation in the absence of a total effect. A further limitation is the relatively low sample size of N = 39. Future work in a larger sample is warranted.

Overall, our study sheds valuable light on processes of emotional learning, on how information encoding and sleep processes may be related, and on differences in these processes in those with and without PTSD and in female and male subjects. With respect to PTSD, we see that failures to differentially tag negative from neutral stimuli at encoding may create a barrier to an effective consolidation process during sleep. Future studies of overnight sleep may show that PTSD group differences in sleep architecture further impair adaptive consolidation processes. As this and other studies highlight, powering studies to properly examine sex differences in these processes is also critical.

## Acknowledgements

This work was supported by the U.S. Department of Veterans Affairs through a VA Career Development Award (Grant No. 5IK2CX000871−05 [to AR]).

