## supplement for "Sleep and emotional memory: translating neural response at encoding to memory accuracy in men and women with and without PTSD"

### Supplementary Methods

#### GAMM Methods

GAMMs are semiparametric statistical models that allow for modelling non-linear relationships in a nonparametric manner while accounting for repeated measurements made on the same subjects and stimuli by incorporating random effects in the models (Wood, 2017). GAMMs were fit using the *bam()* function in the mgcv R package (Wood, 2003, 2004, 2011, 2017; Wood et al., 2016) with a scaled t-distribution with 6.5 degrees of freedom to account for the heavy tails that were seen in QQ-plots of the model residuals while using a normal distribution. The degrees of freedom were chosen based on visual inspection of the QQ-plot of the model residuals. Models also included an autoregressive-1 term with $\rho = 0.85$ to account for autocorrelation in the residuals, with $\rho$ chosen based on the autocorrelation function of the model residuals.

An initial model of the overall ERPs included smooth functions of time (with a cubic spline basis function of dimension k = 40), a smooth factor interaction between stimulus type and time (with a sum-to-zero basis function with k = 40), random smooths for participants (using a factor smooth basis function with k = 40 and m = 1), random intercepts for stimuli, and random intercepts for the stimuli. A next model which allowed for electrode channel differences in the ERP included smooth factor interaction terms between channel and time (sum-to-zero basis function with k = 40) and channel differences in condition were assessed by incorporating a smooth interaction between channel, condition, and time (sum-to-zero basis function with k = 40). Critically, simultaneous confidence regions were computed which ensured that the type-I error rate was controlled across the entire ERP waveform—allowing us to make significance claims at the nominal level regardless of when we observed significant differences in the ERPs.

### Supplementary Results

#### Sex Moderated Mediation

Sex did not significantly moderate the LPP ($b = 0.41$ (0.41), $t(21764) = 1.0$, $p = 0.32$) but sex again moderated the direct effect ($b = 0.65$ (0.10), $z = 6.76$, $p < 0.001$). That is, there were no significant sex differences in the late LPP and there was a significant indirect effect of stimulus type on d' though the late LPP for both women (*indirect effect* = 0.017, 95% bootstrapped CI [0.005, 0.030], $p = 0.006$) and men (*indirect effect* = 0.019, 95% bootstrapped CI [0.005, 0.034], $p = 0.006$), and this indirect effect did not differ between sexes (*Male - Female indirect effect* = 0.002, 95% bootstrapped CI [-0.002, 0.007], $p = 0.32$). The direct effect again differed between sexes with women showing lower d' for negative compared to neutral stimuli (*direct effect* = -0.55, 95% bootstrapped CI [-0.78, -0.32] $p < 0.001$) and men showing no difference (*direct effect* = 0.09, 95% bootstrapped CI [-0.13, 0.31] $p = 0.45$). Women had an overall lower d' for negative compared to neutral stimuli (*total effect* = -0.53, 95% bootstrapped CI: [-0.75, -0.31], $p < 0.001$) and men showed no difference (*total effect* = 0.11, 95% bootstrapped CI [-0.12, 0.33], $p = 0.35$).

#### PTSD Moderated Mediation

The late LPP mediating an emotional memory benefit in both PTSD- (*indirect effect* = 0.020, 95% bootstrapped CI [0.006, 0.036], $p = 0.0052$) and PTSD+ (*indirect effect* = 0.012, 95% bootstrapped CI [0.003, 0.024], $p = 0.0052$) individuals (Figure 9b), however the magnitude of the indirect effect was smaller in PTSD+ individuals (PTSD+ - PTSD- indirect effect = -0.01, 95% bootstrapped CI [-0.017, - 0.002], *p* = 0.005).

**Supplementary Table and Figures**

| Parameter | EDF | Ref. DF | F | p-value |
|---|---|---|---|---|
| **s(time)** | 37.85 | 38.20 | 27.43 | < 0.001 |
| **s(condition, time)** | 35.64 | 39.13 | 45.73 | < 0.001 |
| **s(channel, time)** | 41.88 | 44.90 | 40.84 | < 0.001 |
| **s(channel, condition, time)** | 6.83 | 7.34 | 1.44 | 0.18 |

Supplementary Table 1. Generalize additive mixed effects model parameters. s() refers to smooth function, EDF = effective degrees of freedom. Ref.DF = reference degrees of freedom.

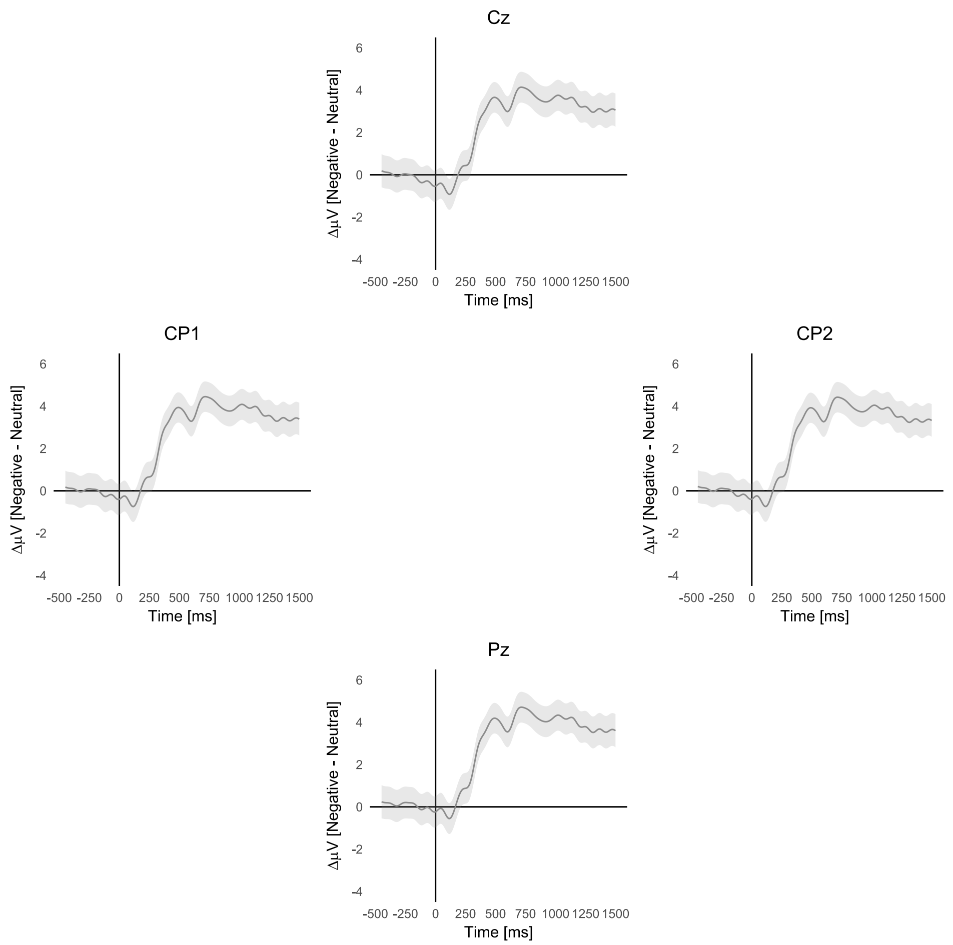


Supplementary Figure 1. GAMM results of the LPP at each of the electrode site.

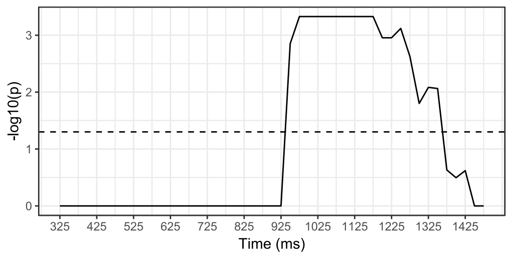


Supplementary Figure 2. -log10(P-values) for the threshold-free cluster enhanced Z-statistics testing the ERP * sleep composite score interaction when predicting d'. P-values above the dotted line are significant adjusting for multiple comparisons across the entire LPP.
